# A fluorescent reporter model for the visualization and characterization of T_DC_

**DOI:** 10.1101/2023.04.06.535573

**Authors:** Alessandra Fiore, Eleonora Sala, Chiara Laura, Michela Riba, Maria Nelli, Valeria Fumagalli, Federico Oberrauch, Marta Mangione, Claudia Cristofani, Paolo Provero, Matteo Iannacone, Mirela Kuka

## Abstract

T_DC_ are hematopoietic cells that combine dendritic cell (DC) and conventional T cell markers and functional properties. They were identified in secondary lymphoid organs (SLOs) of naïve mice as cells expressing CD11c, major histocompatibility molecule (MHC)-II, and the T cell receptor (TCR) β chain. Despite thorough characterization as to their potential functional properties, a physiological role for T_DC_ remains to be determined. Unfortunately, using CD11c as a marker for T_DC_ has the caveat of its upregulation on different cells, including T cells, upon activation. Therefore, a more specific marker is needed to further investigate T_DC_ functions in peripheral organs in different pathological settings. Here we took advantage of Zbtb46-GFP reporter mice to explore the frequency and localization of T_DC_ in peripheral tissues at steady state and upon viral infection. RNA sequencing analysis confirmed that T_DC_ identified with this reporter model have a gene signature that is distinct from conventional T cells and DC. In addition, frequency and total numbers of T_DC_ in the SLOs recapitulated those found using CD11c as a marker. This reporter model allowed for identification of T_DC_ in situ not only in SLOs but also in the liver and lung of naïve mice. Interestingly, we found that T_DC_ numbers in the SLOs increased upon viral infection, suggesting that T_DC_ might play a role during viral infections. In conclusion, we propose a visualization strategy that might shed light on the physiological role of T_DC_ in several pathological contexts, including infection and cancer.

## Introduction

T_DC_ are hematopoietic cells that combine dendritic cell (DC) and conventional T cell markers and functional properties. They were identified by Kuka et al in the SLOs of naïve mice as cells expressing CD11c and MHC-II, two molecules used to identify murine DC, as well as TCRβ, a defining marker for conventional T cells (Kuka et al., 2012; Kuka and Ashwell, 2013). From an ontogenic point of view, T_DC_ are thymus-derived since they were not detected in the spleens of athymic mice and they require the same thymic positive selection as conventional T cells. They are positive for other T cell markers such as CD3, Thy-1, CD27, express CD4 or CD8β at the same ratio as conventional T cells, and present a polyclonal Vβ repertoire comparable to conventional αβ T cells. At steady-state T_DC_ do not display signs of recent activation (CD69, CD25, or IL-7Rhi) or T cell memory markers, excluding the hypothesis that T_DC_ might represent a subsect of activated conventional T cells. In addition, this population does not express other lineage markers, excluding that they might belong to other innate cell subsets such as pDC, NK or NKT (Kuka et al., 2012).

DC are innate cells that bridge innate and adaptive immunity, given their key role in T cell activation. They rely on the expression of MHC molecules and of costimulatory ligands for their antigen-presenting and T cell priming activities (Eisenbarth, 2019). T_DC_ resemble DC in their capacity to expand after FLT3L-mediated stimulation and in the ability to respond to TLR agonists such as LPS. Notably, after TLR stimulation, T_DC_ release IL-12, a cytokine normally produced by DC and important for Th1 polarization. T_DC_ also express CD80/CD86 and *in vitro* studies showed that they can present antigen to CD4^+^ T through MHC-II (Kuka et al., 2012). On the other hand, the TCR expressed by T_DC_ is functional and it can be triggered *in vitro* both by monoclonal antibodies directed to CD3 and by cognate antigens. These experiments also led to the intriguing hypothesis that T_DC_ might be self-sufficient in antigen presentation since they can potentially provide co-stimulation to themselves (Kuka et al., 2012). Finally, T_DC_ were found also in humans: approximately 0.2% of peripheral blood lymphocytes (PBL) are CD3^+^ TCRαβ_+_ CD11c^+^ and HLA-DR^+^ (MHC-II) (Kuka et al., 2012).

These features render T_DC_ the first unconventional polyclonal T cell subset that has ever been described, and potentially key in the context of immune responses. Unfortunately, using CD11c as a marker for T_DC_ has the caveat of its upregulation on some T cell populations upon activation (Qualai et al., 2016; Beyer et al., 2005; Huleatt and Lefrançois, 1995; Lin et al., 2003) thus leading to potentially confounding gating strategies. Since, positivity for CD11c can be used for the identification of T_DC_ in steady-state conditions but not in inflammation or other pathological settings, a more specific marker is needed to further investigate T_DC_ functions in peripheral organs or during inflammation.

Since T_DC_ appear to be developmentally related to both conventional T cells (they need a thymus for development) and to classical DC (they express FLT3 and expand upon FLT3L- mediated stimulation), we asked whether we could rely on the DC-restricted transcription factor Zbtb46 to identify T_DC_ in settings of inflammation and infection (Satpathy et al., 2012). Indeed, Zbtb46 is expressed by all subsets of conventional myeloid dendritic cells (DC), including their direct precursors in the bone marrow (Meredith et al., 2012a; Satpathy et al., 2012). Microarray and qPCR analysis revealed that Zbtb46 was also expressed by T_DC_, indicating that they might derive by the same precursors as classical DC (Kuka et al., 2012). Here we exploited the Zbtb46- GFP reporter mouse model to explore the frequency and localization of T_DC_ in peripheral tissues such as the liver, small intestine, and lung. RNA sequencing analysis confirmed that T_DC_ identified with this reporter model have a gene signature that is distinct from conventional T cells and DC. In addition, frequency and total numbers of T_DC_ in SLOs recapitulated those previously found using CD11c. The Zbtb46-GFP reporter model allowed for the identification of T_DC_ in situ not only in SLOs but also in the liver and lung of naïve mice.

## Results

### Zbtb46 is a reliable marker for the identification of T_DC_

We took advantage of Zbtb46-GFP knock-in reporter mice, which express Zbtb46 and GFP in all cells with an activated Zbtb46 promoter (Satpathy et al., 2012). Despite being a DC-specific marker, it was reported that Zbtb46 expression was not required for DC development (Meredith et al., 2012b). Therefore, both heterozygous and homozygous mice can be used to track cells of DC lineage.

Spleens and lymph nodes (LN) of naïve Zbtb46-GFP homozygous mice were analyzed and characterized in order to validate this fluorescent reporter model. We found that the frequency of classical DC (CD11c^+^MHC-II^+^) and of T_DC_ (CD11c^+^MHC-II^+^TCRβ^+^) in reporter mice was comparable to WT mice, confirming that lack of *Zbtb46* does not affect DC development and recruitment to SLOs (Fig. 1A). To understand whether Zbtb46 expression can be used in place of CD11c to identify DC and T_DC_, we gated DC using both markers. We found that whereas about 70-75% of CD11c^+^MHC-II^+^ cells expressed GFP (Fig. 1B), more than 98% of the GFP^+^MHCII^+^ population was CD11c-positive. These findings indicate that Zbtb46 expression is highly specific and can be used as a valid marker in substitution of CD11c. We then analyzed the percentage of T_DC_, defined as cells positive for the TCRβ chain within the population of GFP^+^MHC-II^+^ cells (Fig. 1C). Frequency and total numbers of T_DC_ in spleens of Zbtb46-GFP mice recapitulated those found using CD11c as a marker, thus confirming the validity of our model (Fig. 1D). Of note, dimensions and granularity of T_DC_ were comparable to those of classical DC, as indicated by the analysis of physical parameters such as Forward Scatter Area (left) and Side Scatter Area (Fig. S1). This observation rules out the possibility that T_DC_ might represent T-DC doublets, a hypothesis that was already thoroughly addressed and excluded in the original report on this new cell population (Kuka et al., 2012).

**Figure 1.**
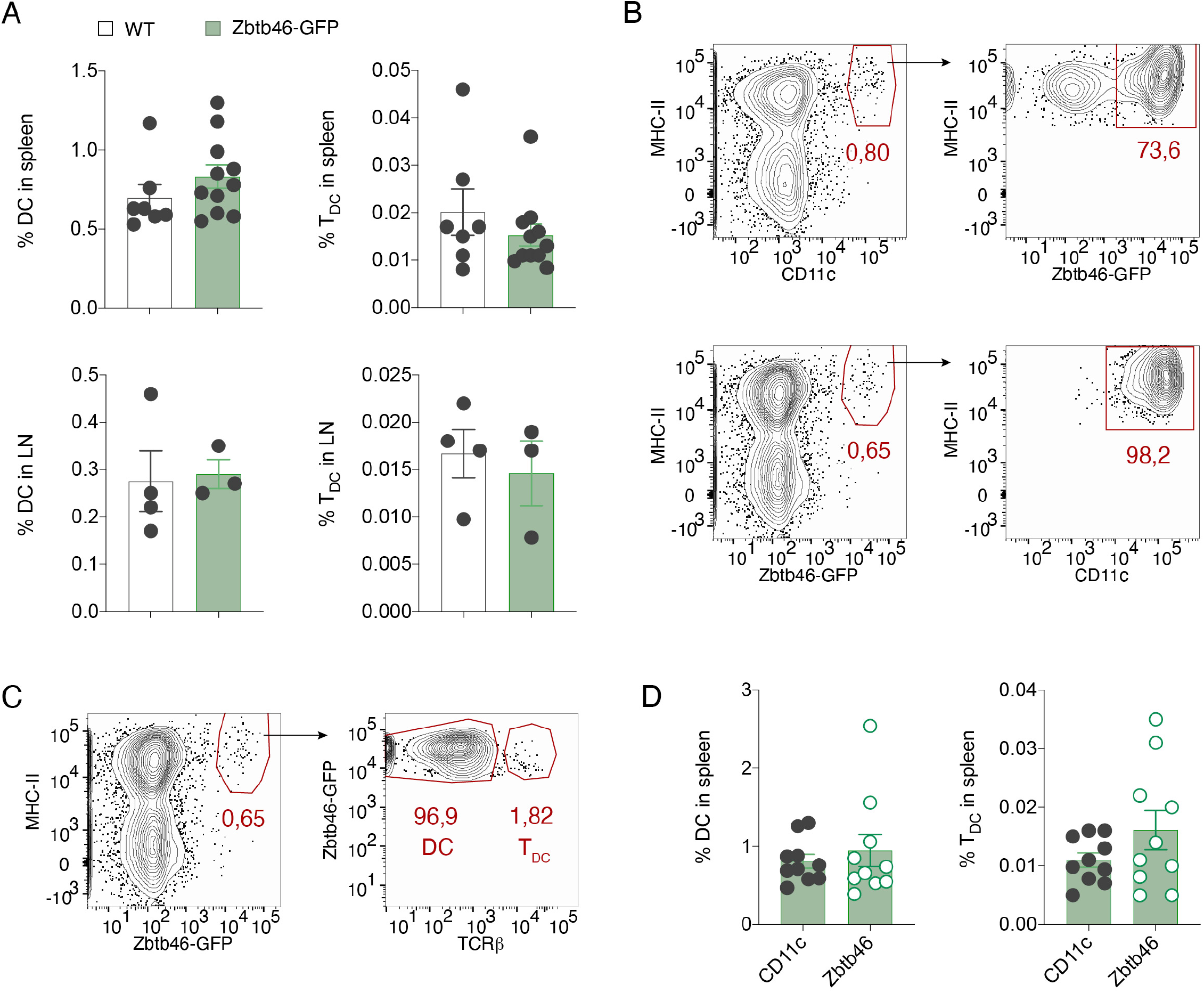
Zbtb46 is a valid substitute of CD11c marker for the identification of T_DC_. **A)** Spleens and popliteal LNs (pLNs) from naive WT or Zbtb46-GFP mice were collected and stained for the typical DC markers, CD11c and MHC-II (I-Ab). Frequencies of DC (left) and T_DC_ (right) in spleens and LNs are shown. *n*=7 (WT spleen), 11 (Zbtb46-GFP spleen), 4 (WT LN), 3 (Zbtb46-GFP LN). Statistics is not shown since there are no statistically significant differences between conditions. **B)** Representative plots showing the frequency of GFP^+^ cells within the CD11c^+^ MHC-II^+^ population (upper panels) or, conversely, plots showing CD11c expression on GFP^+^ MHC-II^+^ cells (lower panels). **C)** Representative plot showing the frequency of DC and T_DC_ within the GFP^+^ MHC-II^+^ population in the spleen of naïve Zbtb46-GFP mice. **D)** Frequencies of DC and T_DC_ gated based on either CD11c or GFP markers in spleens of naïve Zbtb46-GFP mice are shown. *n*=10. Statistics is not shown since there are no statistically significant differences between conditions.

### T_DC_ are distinct from conventional T cells and DC with regard to their transcriptional profile

To investigate the gene expression profile of T_DC_, we performed a bulk RNA sequencing (RNAseq) on T_DC_ from Zbtb46-GFP reporter mice. T_DC_ were sorted from splenocytes of naïve mice along with conventional T cells and DC. The sorting protocol which had been previously optimized for this cell type (Kuka and Ashwell, 2013) yielded almost 100% pure cells (Fig. 2A). Principal component analysis showed that cell identity corresponds to the source of the largest variance in the samples (obtained from three independent experiments) analyzed. This variance is not associated to other characteristics of the set of samples or batch effects (Fig. 2B). In addition, hierarchical clustering confirmed a distinct identity of the three cell types, suggesting however a closer relationship between DC and T_DC_ (Fig. S2A). These data confirm our previous findings that T_DC_ are characterized by a specific cell identity, distinct from conventional T cells and DC (Kuka et al., 2012). 2985 genes were expressed by T_DC_ at higher levels than on conventional T cells (Fig. 2C). Among these genes, we first selected the top 100 differentially expressed genes by T_DC_ with respect to conventional T cells (Fig. 2D). Some of these genes were expressed by both T_DC_ and DC and this strongly suggests that T_DC_ signature in part resembles that of classical DC. We further confirmed this by performing a network analysis on genes that 1) were differentially expressed by T_DC_ versus T cells 2) were expressed at least 50 counts per gene in T_DC_, and 3) were annotated as Zbtb46 neighbours in the STRING protein-protein interaction network database (Fig. S2B) (Szklarczyk et al., 2023). Many genes known to be expressed by the DC lineage (i.e. *Flt3, Spi1, Itgax, Sirpa, IRF8, Batf3, Csf1r*) were found in this network (MacDonald et al., 2005; Mak et al., 2011; Miller et al., 2012). Please note that Zbtb46 itself is not expressed because cells were sorted from homozygous Zbtb46-GFP mice (Fig. S2B). 1647 genes were expressed at significantly higher levels in T_DC_ with respect to DC (Fig. 2C). By performing a Gene Set Enrichment Analysis (GSEA) analysis (Subramanian et al., 2005), we found that the T cell signature extracted from the Panglao database (Franzén et al., 2019) is significantly enriched with some of these genes (Fig. S2C). We then proceeded in selecting the top 100 differentially expressed genes by T_DC_ with respect to classical splenic DC (Fig. 2E). Most of the genes shown in the heatmap of Fig. 2E are specifically expressed by T_DC_ with respect to both conventional T cells and DC, whereas some of them are shared with T cells. Finally, we asked whether T_DC_ sorted from Zbtb46-GFP reporter mice express typical cytotoxic genes, as previously reported for T_DC_ identified via CD11c (Kuka et al., 2012). Although with a high degree of variability among biological replicates, genes like *Gzmb, Gzma, Nkg7, Prf1,* and *Ifng* were expressed at higher levels by T_DC_ with respect to DC (Fig. S2D). Overall, these data confirm that T_DC_ identified with the Zbtb46-GFP reporter mouse are distinct from both conventional T cells as well as DCs.

**Figure 2.**
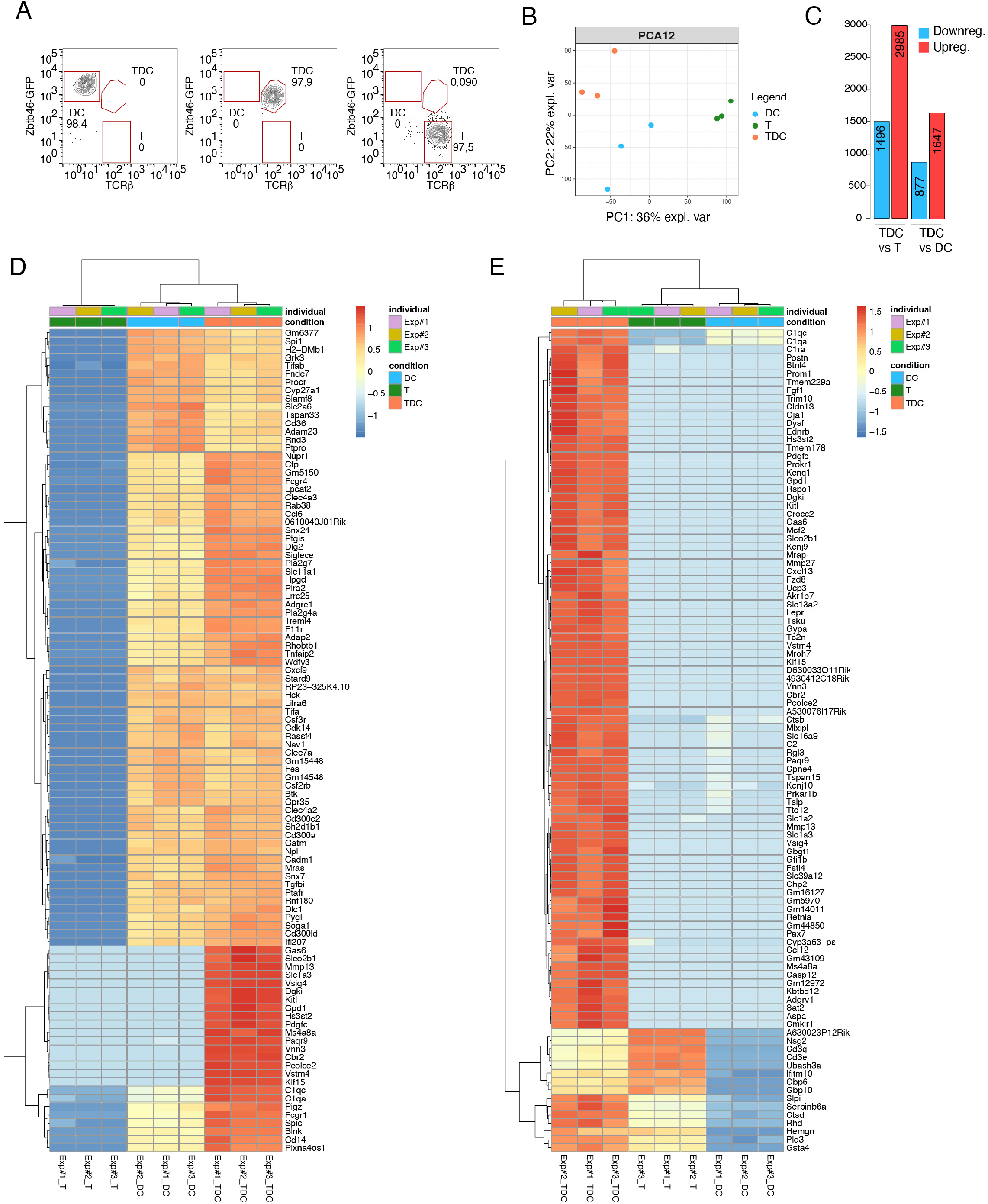
T_DC_ are distinct from conventional T cells and DC with regard to their transcriptional profile. **A)** Splenocytes from 4-5 naïve Zbtb46-GFP mice were pooled and DC, T cells and T_DC_ were sorted. A representative plot of the sorting purity for each cell type is shown. **B)** Representation of the first two components of Principal component analysis (PCA) accounting for the largest variance in the dataset showing separation of samples according to cell type. **C)** Bar plot showing the number of downregulated (blue) and upregulated (red) genes in the indicated comparisons and according to the following cut-off: nominal pvalue < 0.01 and |log2fold change| > 1. **D)** Heatmap showing the top 100 differentially expressed genes in T_DC_ versus T cells. Values in log2(RPKM) were scaled by row across samples. **E)** Heatmap showing the top 100 differentially expressed genes in T_DC_ versus DC. Values in log2(RPKM) were scaled by row across samples.

### T_DC_ can be identified in situ in SLOs and peripheral organs of naïve Zbtb46-GFP bone marrow (BM) chimeras

We asked if we could identify T_DC_ in situ in SLOs from naïve mice by using the Zbtb46-GFP reporter. Although the original paper describing the Zbtb46-GFP mouse model reported GFP expression in a small fraction of endothelial cells (Satpathy et al., 2012), we found a much higher expression than expected, at the point that we could not distinguish DC from stromal cells in SLOs (Fig. S3A and B). Indeed most of the GFP signal was located close to the LN subcapsular sinus (denoted by the CD169 staining) and outside the splenic white pulp (also denoted by the CD169 staining), respectively, instead of being localized to the LN paracortex or to the white pulp where DC reside (Eisenbarth, 2019). Therefore, we decided to generate BM chimeras to restrict GFP expression to the hematopoietic compartment. This allowed for the GFP signal to be detected exclusively in DCs and localized to the LN paracortex and to the splenic white pulp (Fig. S3C and D). We then looked for T_DC_ in situ by staining sections from Zbtb46-GFP BM chimeras with an anti-TCRβ antibody. Confocal microscopy in LN sections revealed rare cells positive for GFP and TCRβ concomitantly (Fig. 3A). These cells were frequently located in interfollicular areas, close to B cell follicles or to the subcapsular sinus (Fig. S4A). Similar cells were also detected in the white pulp of spleens from naïve reporter BM chimeras (Fig. 3B and S4B). Interestingly, confocal imaging of a Granzyme B (GzmB)-Tomato fluorescent reporter model (Mouchacca et al., 2013) showed that some GFP^+^ TCRβ_+_ cells also expressed GzmB (Fig. 3C), in line with previously published data showing T_DC_ as having a cytotoxic profile (Kuka et al., 2012). However, due to the very low frequency of GzmB positivity amongst GFP^+^TCRβ_+_ cells, we decided to continue our characterization of T_DC_ with the Zbtb46-GFP reporter only, combined with TCRβ staining.

**Figure 3.**
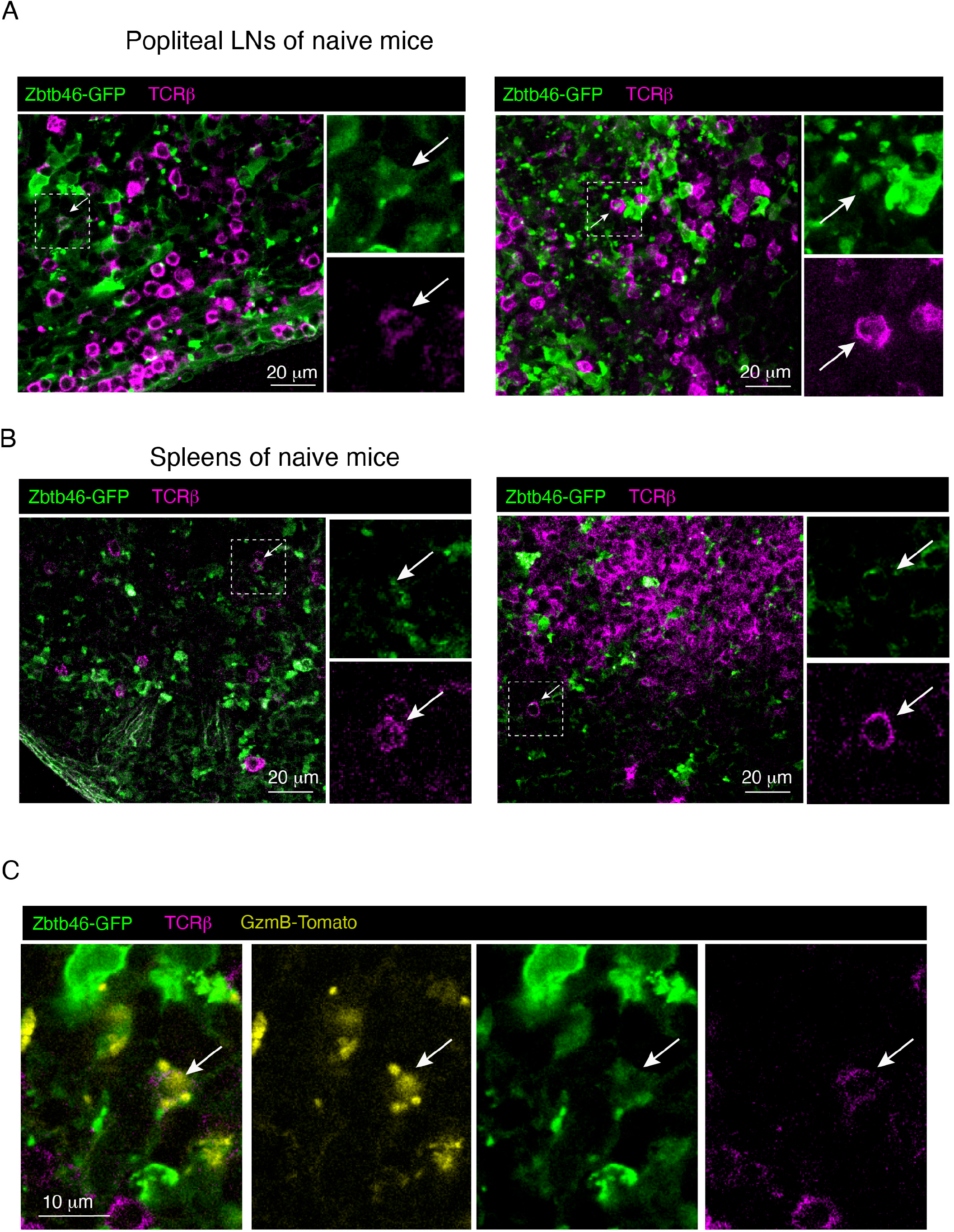
T_DC_ can be identified in situ in SLOs of naïve Zbtb46-reporter BM chimeras. **A-B)** pLNs (A) and spleens (B) from naive Zbtb46-GFP BM chimeras were analyzed by confocal microscopy. Confocal micrographs of two representative mice are shown. Scale bars represent 20 μm. Zbtb46-GFP^+^ cells are in green, TCRβ_+_ cells are in purple. Cells that express both markers concomitantly are indicated with an arrow, and a magnification is shown on the right. **C)** A representative confocal micrograph of a naïve LN showing a cell that expresses concomitantly Zbtb46-GFP, TCRβ, and GzmB-Tomato. The scale bar represents 10 μm.

As other innate lymphocytes or lymphoid cells, T_DC_ might locate not only to SLOs but also to peripheral tissues, so that they can respond to infections in a prompt way. We decided to exploit the Zbtb46-GFP reporter model to explore the presence of T_DC_ in peripheral tissues often in contact with pathogens. We started with the lung, since this is a very important infection site for all respiratory viruses (Chiu and Openshaw, 2015). Confocal imaging of perfused lung sections contained rare cells expressing both GFP and TCRβ, suggesting that the lung might be a localization site of T_DC_ at steady state (Fig. 4A). Flow cytometry analysis showed that the frequency of T_DC_ in the lungs was comparable to that of the spleens of the same animals (Fig. 4C). Next, we looked at the liver, which is in contact with many antigens derived from the gut as well as being an infection site for both hepatotropic and systemic viruses (Heymann and Tacke, 2016; Ficht and Iannacone, 2020; Zinkernagel et al., 1986). Cells expressing GFP and TCRβ were identified in the parenchyma of perfused livers from naïve mice (Fig. 4B); interestingly, the frequency of T_DC_ (detected by flow cytometry) among intrahepatic leucocytes (IHL) was significantly higher than in the spleens of the same mice (Fig. 4D). These findings indicate that the liver might be a preferential location site for T_DC_. By contrast, we found a very low frequency of Zbtb46-positive cells in the small intestine, and they were almost all DC, with very few and almost undetectable T_DC_, suggesting that this organ might not be populated by T_DC_ at steady state conditions (Fig. 4E).

**Figure 4.**
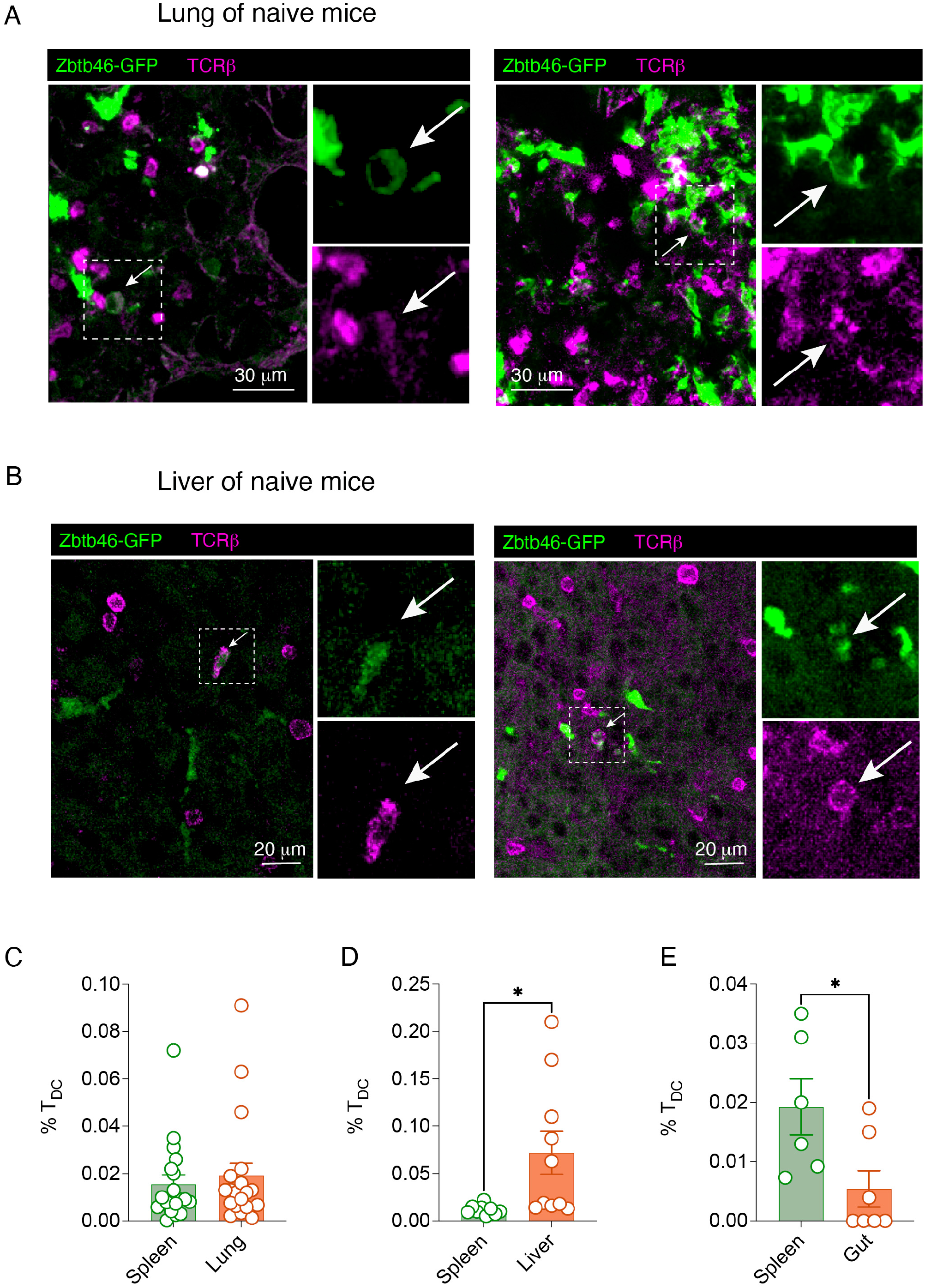
T_DC_ can be identified in lungs and livers of naïve Zbtb46-reporter mice. **A-B)** Lungs (A) and livers (B) from naive Zbtb46-GFP BM chimeras were analyzed by confocal microscopy. Confocal micrographs of two representative mice are shown. Scale bars represent 30 μm (A) and 20 μm (B). Zbtb46-GFP^+^ cells are in green, TCRβ_+_ cells are in purple. Cells that express both markers concomitantly are indicated with an arrow, and a magnification is shown on the right. **C-E)** Frequencies of T_DC_ in lungs (C), livers (D) and gut (E) of naïve Zbtb46-GFP mice analyzed by flow cytometry are shown. *n*=19 (spleen vs lung), *n*=10 (spleen vs liver), *n*=6 (spleen vs gut). *\* p value* < 0.05.

### Viral infection leads to enrichment of T_DC_ in the SLOs

We previously showed that adoptively transferred Ag-specific T_DC_ could expand in response to systemic infection with lymphocytic choriomeningitis virus (LCMV) (Kuka et al., 2012). However, because CD11c is upregulated on some T cell populations during immune activation (Qualai et al., 2016; Beyer et al., 2005; Huleatt and Lefrançois, 1995; Lin et al., 2003), it is difficult to establish whether numbers and frequencies of endogenous T_DC_ change during viral infection. In particular with regard to LCMV infection, the gating strategy based on CD11c and MHC-II double positive cells is not ideal for T_DC_ identification, since at day seven post infection the frequency of CD11c^+^MHC-II^+^ cells is substantially increased (Fig. S5A) mainly due to the fact that the majority of TCRβ_+_ cells acquire CD11c expression (Fig. S5B). Importantly, Zbtb46 expression profile did not change upon LCMV infection (Fig. S5C) and thus the Zbtb46-GFP reporter represents an ideal model to investigate whether T_DC_ expand in LCMV-infected mice. LCMV infection resulted in a significant increase in the frequency of both T_DC_ and conventional T cells in the spleens of LCMV-infected mice analyzed seven days after infection (Fig. 5A). This effect was specific to these two cell types, as frequencies of classical DC did not change significantly (if anything they were found in lower frequency in LCMV-infected mice) (Fig. 5A). Mice infected subcutaneously (s.c.) in the footpad showed a slightly different trend (Fig. 5B): whereas a higher frequency of DC was found in the draining LNs (dLNs) seven days post infection (Fig. 5B), the frequency of total T cells and total T_DC_ did not increase like in the spleen (Fig. S6). We reasoned that, since LCMV is known to strongly expand CD8^+^ T cells, the total numer of T cells upon s.c. infection might reflect the aftermath of a high increase in the CD8 T cell compartment and a relative decrease in the CD4 T cell compartment. Indeed, when we analyzed only CD8^+^ T cells and CD8^+^T_DC_, we could appreciate a substantial increase in both cell populations upon infection (Fig. 5B). Confocal imaging of LN sections from infected mice presented many cells positive for both GFP and TCRβ, thus confirming an enrichment of T_DC_ in these organs upon infection (Fig. 5C). Taken together, these results indicate that T_DC_ expand in SLOs in response to LCMV infection.

**Figure 5.**
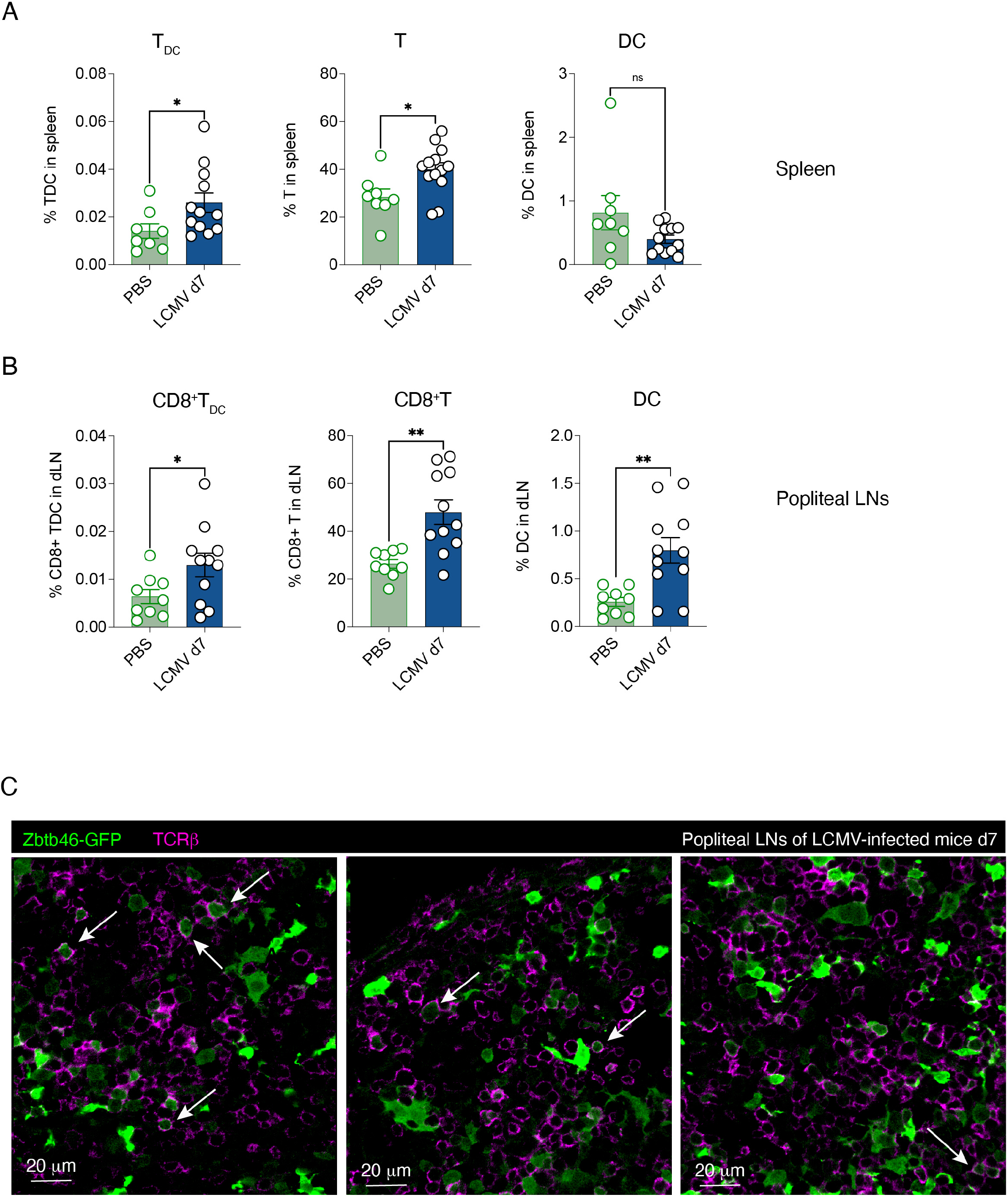
Viral infection leads to enrichment of T_DC_ in the SLOs. Zbtb46-GFP mice (**A-B**) or BM chimeras (**C**) were infected i.v. or s.c. with LCMV Arm, and SLOs were analyzed seven days upon infection. **A)** Frequencies of total T_DC,_ conventional T cells, and DC in the spleens of i.v. infected mice are shown. *n*=8 (PBS), *n*=12 (LCMV). *\* p value* < 0.05. **B)** Frequencies of CD8^+^ T_DC,_ CD8^+^ T cells, and total DC in the draining LNs (dLN) of s.c. infected mice are shown. *n*=9 (PBS), *n*=11 (LCMV). *\* p value* < 0.05*, ** p value* < 0.01. **C)** dLNs from LCMV-infected Zbtb46-GFP BM chimeras were analyzed by confocal microscopy seven days upon infection. Confocal micrographs of three representative sections are shown. Scale bars represent 20 μm. Zbtb46-GFP^+^ cells are in green, TCRβ_+_ cells are in purple. Cells that express both markers concomitantly are indicated with an arrow.

Since T_DC_ represent a population of T cells with innate traits, we asked whether T_DC_ might start expanding earlier than conventional T cells upon infection. Two days upon systemic LCMV the frequency of T_DC_ in the spleens of infected mice was higher than non-infected controls, although the difference was not statistically significant (Fig. S7A). A similar trend was observed for CD8^+^ T_DC_ in the dLNs of subcutaneously-infected mice, whereas no changes were observed in total T_DC_ (Fig. S7B and C). The frequency of conventional T cells and DC did not increase at this timepoint, suggesting a specific and distinct dynamics for T_DC_ that requires further investigation.

### The transcriptional profile of T_DC_ overlaps with that of MyT, a subset of T cells found in the gut during bacterial infection

Recently, a new subset of CD4^+^ T cells was identified in the gut of mice infected with *Salmonella* (Kiner et al., 2021). These cells were named MyT since they expressed both T cell markers and myeloid cell markers. We asked whether MyT and T_DC_ might represent the same cell type, and to test this hypothesis we decided to compare these two populations at the transcriptomic level. First, we re-analyzed the published scRNAseq dataset reporting the existence of MyT (Kiner et al., 2021), focusing on T_eff_ cells only as the authors did. We performed dimensionality reduction on t-distributed stochastic neighbor embedding (t-SNE) plot, highlighting the pathogen used to infect the mice (Fig. 6A). Then, MyT cells were identified among the cells belonging to the *Salmonella* infection condition (Fig. 6B). This annotation was based on the expression of both myeloid cell markers (*H2-Ab1*, *C1qa*, *Lyz2*, *Apoe*) and T cell markers (*Cd3d* and *Trac*), as previously reported (Kiner et al., 2021) and as shown in Fig. S8. We then compared the transcriptome of these cells with the one of T_DC_ obtained from the bulk RNA sequencing experiment previously performed (Fig. 2). To this end, we built a T_DC_ signature using the top 100 differentially expressed genes by T_DC_ with respect to conventional T cells in the bulk RNA sequencing experiment (Fig. 2D). We observed a significant enrichment of this signature in the MyT population compared to the other T cells (Fig. 6C and D). Moreover, the genes belonging to the previously published T_DC_ signature obtained as the ones upregulated in T_DC_ vs T cells (Kuka et al., 2012) are also highly expressed by the MyT population (Fig. S9). Overall, these data strongly suggest that T_DC_ and MyT might represent similar if not identical cell populations.

**Figure 6.**
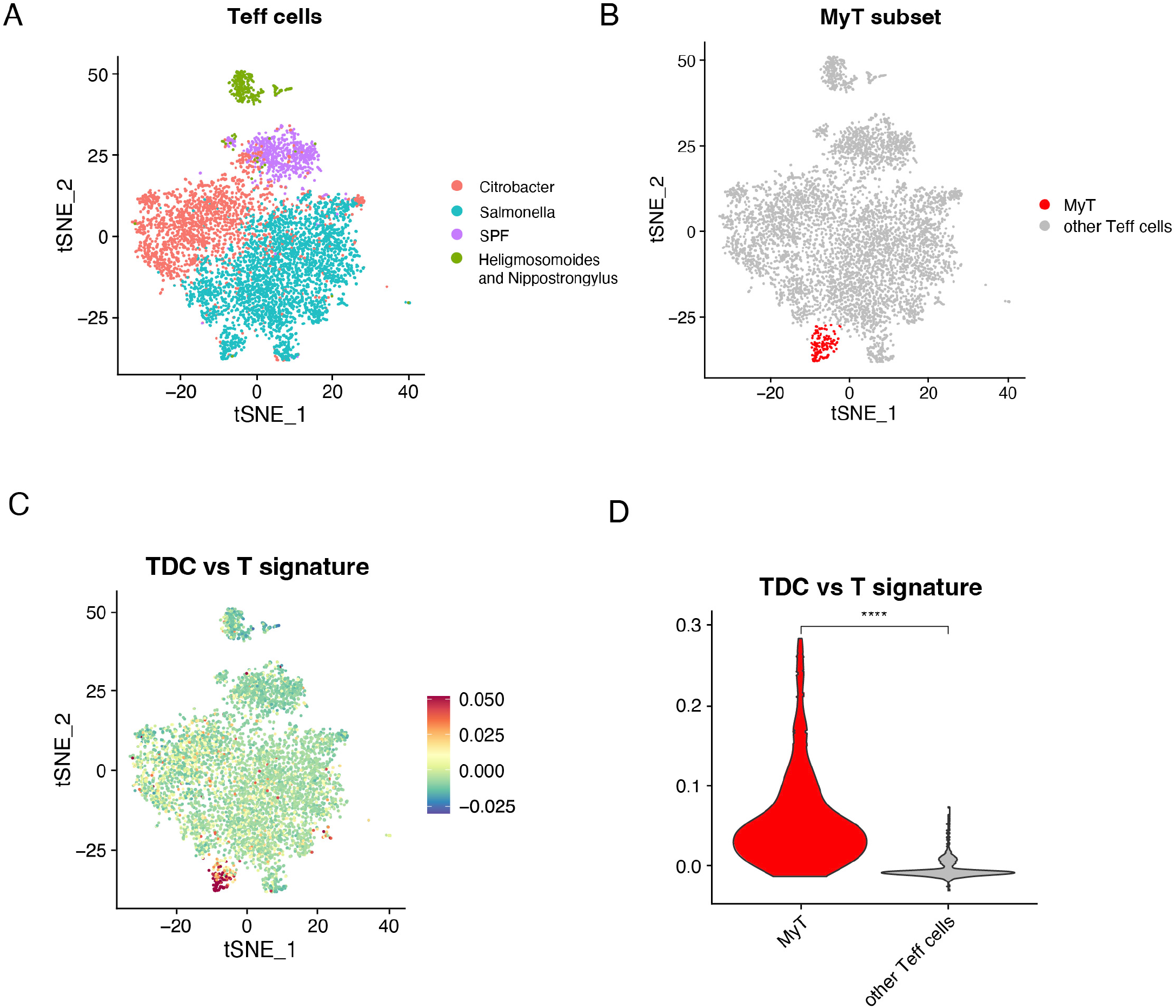
The transcriptional profile of T_DC_ overlaps with that of MyT, a subset of T cells found in the gut during bacterial infection. **A)** t-SNE representation of effector T cells from Kiner et al. Each dot corresponds to a single cell, colored according to the infectious conditions. **B**) Highlight of the MyT population in the t-SNE in (A) based on the expression of genes in figure S8. (**C**) Feature plot of a T_DC_ signature (top 100 DEGs in the comparison TDC vs T) obtained from the bulk RNA seq experiment, max.cutoff parameter set to “q95”. (**D**) Violin plot showing the enrichment of the T_DC_ signature in the MyT population compared to the other T cells. \*\*\*\**p value* <= 0.0001 using a Wilcoxon test.

## Discussion

T_DC_ are unconventional polyclonal T cells that combine innate and adaptive cell properties. When triggered *in vitro*, they can respond either as DC or as conventional T cells, depending on the stimulus (Kuka et al., 2012). However, these represent only potential functional properties of T_DC_ and there is no formal proof that their behavior *in vivo* is similar. In general, the physiological role and relevance of T_DC_ *in vivo* is unknown. The main reason for this is a technical challenge in precisely identifying them due to the low frequency of these cells and the lack of specific markers that can be exploited to generate T_DC_-deficient mice. In addition, the combination of markers used to identify T_DC_ in the SLOs of naïve mice are not ideal for their identification in peripheral organs or during inflammation and infection, conditions where the integrin CD11c is widely expressed also in cell types other than the DC lineage (Qualai et al., 2016; Beyer et al., 2005; Huleatt and Lefrançois, 1995; Lin et al., 2003). Here we tackle this last caveat by taking advantage of a fluorescent reporter mouse strain already used in the past to identify DC (Meredith et al., 2012a; Satpathy et al., 2012; Meredith et al., 2012b). Thanks to this reporter model we could detect T_DC_ in situ in SLOs, lung, liver, both at steady state and upon viral infection.

In the SLOs of naïve mice T_DC_ cells were frequently located in interfollicular areas, close to B cell follicles or to the subcapsular sinus, reminiscent of a previously described innate αβ CD8 T cell population previously described by Kastenmuller *et al* (Kastenmuller et al., 2012). These innate αβ CD8 T cells were found to be among the main producers of IFNγ very early after infection and located to the subcapsular sinus, but were not further characterized. Of note, bulk RNA sequencing showed that T_DC_ of naïve mice are highly enriched in *Ifng* expression (Fig. S1D), therefore suggesting that they might overlap with the innate cells described previously (Kastenmuller et al., 2012).

Recently, a subset of intestinal ILC3 expressing Zbtb46 was described (Zhou et al., 2022). Zbtb46 expressed by ILC3 restrains the inflammatory potential of these cells. Up to this report Zbtb46 was known to be expressed only on cells of the DC lineage and on endothelial cells. These latest findings suggest that Zbtb46 is not a DC-restricted marker but it might be expressed also by cells of lymphoid origin. Whereas the prevailing dogma attributes to DC and conventional T cells extremely divergent pathways of differentiation, the identification of genes expressed by both myeloid and lymphoid precursors support a common DC/T ontogeny for T_DC_. For example, IRF8 has been found to be expressed by both T and DC precursors in human thymus, before their commitment to one of the cell lineages (Liang et al., 2023). Notably *Irf8* is among the genes which are differentially expressed by T_DC_ with respect to conventional T cells (Fig. S2B).

The Zbtb46-GFP reporter model led to the finding that, at steady state, T_DC_ preferentially locate to the liver and to a lower extent to the lung, but are excluded by the small intestine. This finding seems in stark contrast with the observation that MyT, a new subset of T cells with myeloid properties and whose transcriptional profile overlaps with the one of T_DC,_ was discovered in the gut in response to bacterial infection (Kiner et al., 2021). One explanation for this controversy might be that the gut is not a preferential site of location in steady-state conditions, but T_DC_ or MyT might migrate there in response to the infection from elsewhere (i.e. from SLOs). Moreover, we found that LCMV infection leads to preferential expansion of CD8^+^ T_DC_, whereas MyT found during bacterial infection express CD4. We have previously reported that in naïve mice T_DC_ can express either CD4 or CD8. This would suggest that different subsets of T_DC_ might be expanded upon infection, depending on the pathogen. Pathogens that lead to strong CD8 T cell responses like LCMV might lead to expansion of CD8^+^ T_DC_, whereas pathogens that trigger CD4 T cell responses might lead to expansion of CD4^+^ T_DC_.

A recent study shows that the liver is populated by CD8^+^ T cells with myeloid markers on their surface (Pallett et al., 2023). The authors showed that these myeloid markers (among which CD14) are acquired by T cells following their activation by myeloid cells and result in functional changes in T cell function. However these hybrid CD8^+^ T cells do not express any RNA related to those myeloid markers. By contrast, both T_DC_ and MyT are characterized by a transcriptional profile that combines both myeloid and T cell markers, therefore we strongly believe that the hepatic CD14^+^CD8^+^ T cells are different from T_DC_.

In conclusion, we believe that the T_DC_ visualization strategy we propose might shed light on the physiological role of T_DC_ in several pathological contexts, including infection and cancer. Indeed, further investigation on the localization of T_DC_ in specific organ sub-compartments and of their interactions with other cell types, as well as on their enrichment upon specific conditions, might provide new information that can partially overcome the caveat of the temporary lack of a specific marker that could be used for more functional studies.

## Materials and Methods

### Mice

Mice were housed under specific pathogen-free conditions and used at 8-10 weeks of age, unless otherwise indicated. All experimental animal procedures were approved by the Institutional Animal Committee of the San Raffaele Scientific Institute.

B6.129S6(C)-Zbtb46tm1.1Kmm/J mice (in the text referred to as Zbtb46-GFP) were purchased from The Jackson Laboratory. C57BL/6 were purchased from Charles River. GzmB-Tomato mice were kindly provided by Dr. Claude Boyer (Mouchacca et al., 2012; Mouchacca et al., 2013). Bone marrow chimeras were generated by irradiation of C57BL/6 mice with ∼900 rad and reconstitution with the indicated bone marrow; mice were allowed to reconstitute for at least 8 weeks prior to use.

### Infections and immunizations

Mice were infected subcutaneously (s.c.) in the footpad with 1x10^5^ focus forming units (ffu) or intravenously (i.v.) with 2x10^5^ ffu of LCMV Armstrong (LCMV-Arm). Virus was propagated and quantified as described (Welsh and Seedhom, 2008) and diluted in 25 µl of PBS prior to s.c. injection or in 200 µl of PBS prior to i.v. injection. All infectious work was performed in designated Biosafety Level 2 (BSL-2) and BSL-3 workspaces in accordance with institutional guidelines.

### Cell Isolation and Flow Cytometry

Single-cell suspensions of spleens and LNs were generated as described (Sammicheli et al., 2016). For lungs analysis, mice were perfused through the right ventricle with PBS. Lung tissue was digested in RPMI 1640 containing 3.2 mg/ml Collagenase IV (Sigma, #C5138) and 25 U/ml DNAse I (Sigma, #D4263) for 30 minutes at 37°C. Homogenized lungs were passed through 70 µm nylon meshes to obtain a single cell suspension. For liver analysis, mice were perfused through the vena cava with PBS. Liver tissues were disrupt using scissors on 70 µm nylon meshes and were digested in RPMI 1640 containing 0.2 mg/ml Collagenase IV and 5 U/ml DNAse I for 40 minutes at 37°C. After, cell suspensions were centrifuged at 300 rpm for 3 minutes and surnatants were recovered. Small intestine was harvested paying attention to remove fat and Peyer patches. It was cut longitudinally and rinsed with PBS, then it was placed in complete medium (DMEM supplemented with 10% FBS, 1% penicillin plus streptomycin, 1% L-glutamine) with 1mM DTT (Sigma, # 10197777001) for 10 minutes at 37°C. The pieces of small intestine were then transferred in complete medium with 1mM EDTA for 10 minutes at 37°C. After that, EDTA buffer was replaced with a fresh one for other 10 minutes at 37°C. Tissue suspension was placed in fresh medium with 1 mg/ml Collagenase D (Sigma, # 11088858001) and 5 U/ml DNase I for 30 minutes at 37°C. Homogenized intestine was passed through 70 µm strainer and washed one time with PBS.

Cell suspensions obtained from lung, liver and intestine processing were resuspended with a solution composed by 36% percoll (Sigma #P4937) and 4% PBS 10x in PBS. After centrifugation for 20 minutes at 2000 rpm (light acceleration and low brake), cells were isolated and counted. Possibly, the remaining red blood cells were removed with ACK lysis.

All flow cytometry stainings were performed as described (Kuka et al., 2012; Sammicheli et al., 2016). Antibodies used included: CD4 (RM4-5), TCRβ (H57-597), CD8 (K53-6.7), CD11c (N418), MHCII (AF6-120.1). Fluorochrome-conjugated Abs were purchased from BioLegend, eBioscience or BD Pharmingen. All flow cytometry analyses were performed in FACS buffer containing PBS with 2 mM EDTA and 2% FBS on a FACS CANTO (BD Pharmingen) and analysed with FlowJo software (Treestar).

### Confocal immunofluorescence histology

Confocal microscopy analysis of popliteal LNs, spleens, livers and lungs was performed as previously described (De Giovanni et al., 2020; Sammicheli et al., 2016). The following primary Abs were used for staining: rat anti-B220 (RA3-6B2), rabbit anti-GFP (Invitrogen), anti-TCRβ (H57), anti-CD169 (Ser-4). Images were acquired on an inverted Leica microscope (SP8, Leica Microsystems) with a motorized stage for tiled imaging using a HC PL APO CS2 20X objective (NA 0.75). To minimize fluorophore spectral spill over, we used the Leica sequential laser excitation and detection modality. B cell follicles were defined based on the B220 staining

### Cell Sorting and RNA extraction

Splenocytes of naive Zbtb46-GFP mice were processed in order to obtain single cells suspensions and T cells, DCs and T_DC_ were sorted. Briefly, splenocytes were gated for MHC-II and Zbtb46-GFP in order to identify DCs (MHC-II^+^Zbtb46-GFP^+^) and non-DCs (MHC-II^-^Zbtb46-GFP^-^). Gating on the non-DC population, T cells were identified and sorted (Fig. 34F) as CD3ε^+^TCRβ^+^ cells. Gating on the MHC-II^+^Zbtb46-GFP^+^population, classical DCs were identified and sorted as CD3ε^-^TCRβ^-^ cells, whereas T_DC_ were identified and sorted as CD3ε^+^TCRβ^+^ cells. The three cell types underwent two rounds of sorting to obtain a higher purity as described in (Kuka and Ashwell, 2013). Total RNA was isolated with the RNeasy Micro kit (Qiagen) from 2,000 to 3,000 cells and then subjected to bulk RNA sequencing.

### RNA-seq data processing and analysis

Sequencing Libraries were prepared using SMART Nextera unstranded protocol. Libraries were checked using Qubit (fluorimeter) and Bioanalyzer (capillary electrophoresis). Sequencing was performed using Illumina Nextseq 500 with a HighOutput flow cell, 1x75 nt, single read, and Novaseq 6000, 1X 100 nt, single read. Libraries were found to be of good quality.

FastQC software was used to examine quality of fastq files (Andrews, 2010). Raw sequencing files were trimmed to eliminate adapter sequences, and those trimmed sequences were aligned to the ‘mm10’ mouse genome using STAR aligner (version STAR_2.5.3a) (Dobin et al., 2013). (Liao et al., 2019) with the featureCounts function was used for counting the abundance of genes. Principal Component Analysis was performed to evaluate the separation of samples based on decreasing variance. Putative differentially expressed genes were selected using *limma*-voom (Ritchie et al., 2015). The criterion used to select differentially expressed genes in pairwise comparisons is the SEQC cut-off: nominal pvalue <0.01 and absolute value of log2 fold change >1 (Consortium, 2014).

### Single-cell RNA-seq

Raw count datasets from Kiner et al. were downloaded from the Gene Expression Omnibus (GEO) database under accession no. GSE160055. Specifically, the four datasets corresponding to four infection conditions were analysed: GSM4859313_SPF, GSM4859314_Citrobacter, GSM4859315_Salmonella and GSM4859316_Nippo. Single cell data analysis was performed using Seurat (v4.0.1) (Stuart et al., 2019). 6509 cells were obtained after applying the same QC filters used in Kiner et al., i.e. cells with less than 1,000 UMIs or 400 genes and more than 4,000 UMIs or 0.05% of reads mapped to mitochondrial genes were excluded from the analysis.

Moreover, only genes expressed in at least 5 cells were retained. Samples were merged and the UMI count matrix was further normalized and scaled following the standard Seurat workflow. UMAP reduction was then applied on the first 25 Principal Components after running PCA.

The plots showing normalized expression values with a color scale on top of UMAP plots and the Violin plot were produced with FeaturePlot and VlnPlot Seurat functions, respectively. The gene signature average for T_DC_ marker genes was calculated with the AddModuleScore function in Seurat.

### Statistical analyses

Flow and imaging data were collected using FlowJo Version 10.5.3 (Treestar) and Imaris (Bitplane), respectively. Statistical analyses were performed with GraphPad Prism software version 9.5 (GraphPad). Results are expressed as mean ± SEM. Means between two groups were compared with unpaired two-tailed t test. Means among three groups were compared with one-way ANOVA. Tukey’s post-test was used for multiple comparisons. Significance is indicated as follows: * *p value* < 0.05; ** *p value* < 0.01; *** *p value* < 0.001; **** *p value* < 0.0001. Comparisons are not statistically significant unless indicated.

## Acknowledgments

Flow cytometry was carried out at FRACTAL, a flow cytometry resource and advanced cytometry technical applications laboratory established by the San Raffaele Scientific Institute. Confocal immunofluorescence histology was carried out at Alembic, an advanced microscopy laboratory established by the San Raffaele Scientific Institute and the Vita-Salute San Raffaele University. We are grateful to Claude Boyer (Centre d’Immunologie de Marseille-Luminy, Institut National de la Santé et de la Recherche Médicale (INSERM)-Centre National de la Recherche Scientifique (CNRS)-Univ.Med., Parc Scientifique de Luminy) for kindly providing GzmB- Tomato mice. We thank Jonathan D. Ashwell (NCI/NIH, Maryland, US), Anna Mondino, Guido Poli, Paolo Dellabona and Luca G. Guidotti (San Raffaele Scientific Institute, Milan, IT) for helpful discussions. This research was supported by the Italian Ministry of Education, University and Research grant SIR-RBSI14BAO5 to M.K. M.I. is supported by European Research Council (ERC) Consolidator Grant 725038, ERC Proof of Concept Grant 957502, Italian Association for Cancer Research (AIRC) Grants 19891 and 22737, Italian Ministry of Health (MoH) Grant RF- 2018-12365801, Italian Ministry for University and Research (Project no. PE00000007, INF- ACT), and sponsored research agreements from Gilead Sciences, Asher Biotherapeutics and VIR Biotechnology.

## Author contributions

Conceptualization, M.K; Investigation, A.F., E.S., M.N., V.F., F.O., M.M., C.C.; Resources, M.K., M.I; Formal Analysis, M.K, A.F.; Bioinformatic analysis: C.L., M.R., P.P.; Writing M.K. with input from all authors; Visualization, M.K.; Project supervision, M.K.; Funding Acquisition, M.K.

## Data and materials availability

Accession numbers will be made available prior to publication.

## Supplementary Figure Legends

Figure S1. Dimensions and granularity of T_DC_ are comparable to those of DC. Physical parameters of conventional T cells, DC, and T_DC_ from naïve Zbtb46-GFP mice were analyzed and compared through calculation of the geometric mean of the Forward Scatter Area (left) and the Side Scatter Area (right). *n*=4. *\*\* p value* < 0.01; **** *p value* < 0.0001

Figure S2. T_DC_ share expression of some genes with DC and some genes with conventional T cells, but also express a set of cytotoxic genes. **A)** Hierarchical clustering on samples using 16099 expressed genes (at least 5 counts in at least 3 samples) showing separation according to cell types. **B)** STRING database subnetwork representing the first neighbours of Zbtb46 starting from a network obtained selecting from 1538 up-regulated genes in T_DC_ versus T comparison selected excluding genes with less than 50 counts per sample (in T_DC_ group of samples). The procedure was done using cytoscape software (Shannon et al., 2003) and data retrieval was performed with default parameters. The colors reflect the log2 Fold change values in T_DC_ vs T comparison. **C)** GSEA enrichment plot showing up-regulation of the signal of T cell marker gene set downloaded from panglao database (114 genes) (Franzén et al., 2019) in T_DC_ (red) versus DC (blue). **D)** Heatmap showing a set of cytotoxic genes expressed in T_DC_ and in DC. Values in log2(RPKM) were scaled by row across samples.

Figure S3. Zbtb46-GFP BM chimeras allow for specific detection of DC in situ. **A)** A representative confocal micrograph of a pLN from naive Zbtb46-GFP mice is shown. The scale bar represents 200 μm. Zbtb46-GFP^+^ cells are in green, CD169^+^ cells (subcapsular sinus macrophages) are in white. **B)** A representative confocal micrograph of a spleen from naive Zbtb46-GFP mice is shown. The scale bar represents 200 μm. Zbtb46-GFP^+^ cells are in green, CD169^+^ cells (marginal sinus macrophages delimiting the white pulp) are in white. **C)** A representative confocal micrograph of a pLN from naive Zbtb46-GFP BM chimeras is shown. The scale bar represents 150 μm. Zbtb46-GFP^+^ cells are in green, B220^+^ cells (expressed by B cells in the B cell follicles) are in white. **D)** A representative confocal micrograph of a spleen from naive Zbtb46-GFP BM chimeras is shown. The scale bar represents 100 μm. Zbtb46-GFP^+^ cells are in green, B220^+^ cells (expressed by B cells in the B cell follicles) are in white.

Figure S4. T_DC_ are located in interfollicular areas, close to B cell follicles or to the subcapsular sinus. **A-B)** pLNs (A) and spleens (B) from naive Zbtb46-GFP BM chimeras were analyzed by confocal microscopy. Confocal micrographs of two representative mice are shown. Scale bars represent 20μm. Zbtb46-GFP^+^ cells are in green, TCRβ_+_ cells are in purple, B220^+^ cells (expressed by B cells in the B cell follicles) are in white. T_DC_ are indicated with an arrow only in the images where B220 is not shown, for clarity purposes.

Figure S5. Zbtb46, in contrast to CD11c, is not upregulated by T cells upon LCMV infection. **A-C)** Representative plots showing the frequency in the spleen of CD11c^+^MHC-II^+^ cells (A), of CD11c^+^TCRβ_+_ cells (B) or of Zbtb46-GFP^+^MHC-II^+^ cells (C) before and seven days after LCMV infection.

Figure S6. *Frequencies of T_DC_ in SLOs of LCMV-infected mice seven days after infection* Frequencies of total T_DC_ and conventional T cells in the draining LNs (dLN) of s.c. infected mice are shown. *n*=9 (PBS), *n*=11 (LCMV).

Figure S7. Frequencies of T_DC_ in SLOs of LCMV-infected mice early after infection. **A)** Frequencies of total T_DC,_ conventional T cells, and DC in the spleens of i.v. infected mice two days after infection are shown. *n*=6 (PBS), *n*=8 (LCMV). **B)** Frequencies of total T_DC,_ conventional T cells, and DC in the draining LNs (dLN) of s.c. infected mice two days after infection are shown. *n*=4 (PBS), *n*=3 (LCMV). **C)** Frequencies of CD8^+^ T_DC_ and CD8^+^ T cells in the draining LNs (dLN) of s.c. infected mice two days after infection are shown. *n*=4 (PBS), *n*=3 (LCMV).

Figure S8. Markers used for the annotation of the MyT signature **A)** Feature plot representation of the normalized expression level of both myeloid (*Lyz2, Apoe, H2-Ab1, C1qa*) and T cell (*Trac, Cd3d*) markers on the scRNAseq dataset described in Fig. 6, as shown by Kiner et al. (max.cutoff parameter set to “q95” in the FeaturePlot function).

Figure S9. The previously published T_DC_ signature is highly expressed by the MyT population **A)** Feature plot representation on dataset in Fig. 6 of a signature composed by genes in Kuka et al. (Table 1) shown to be upregulated in the comparison T_DC_ vs T cells (max.cutoff parameter set to “q95” in the FeaturePlot function).

