## Supplementary figures and images for "A fluorescent reporter model for the visualization and characterization of T_DC_"

### Supplementary Figures

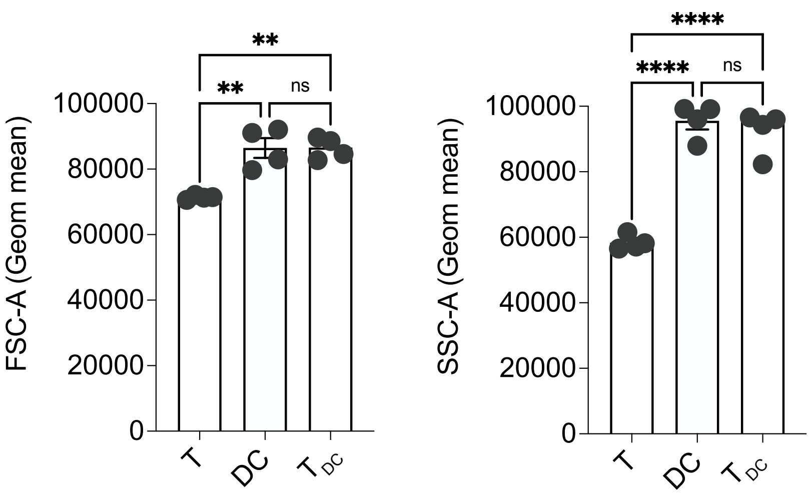

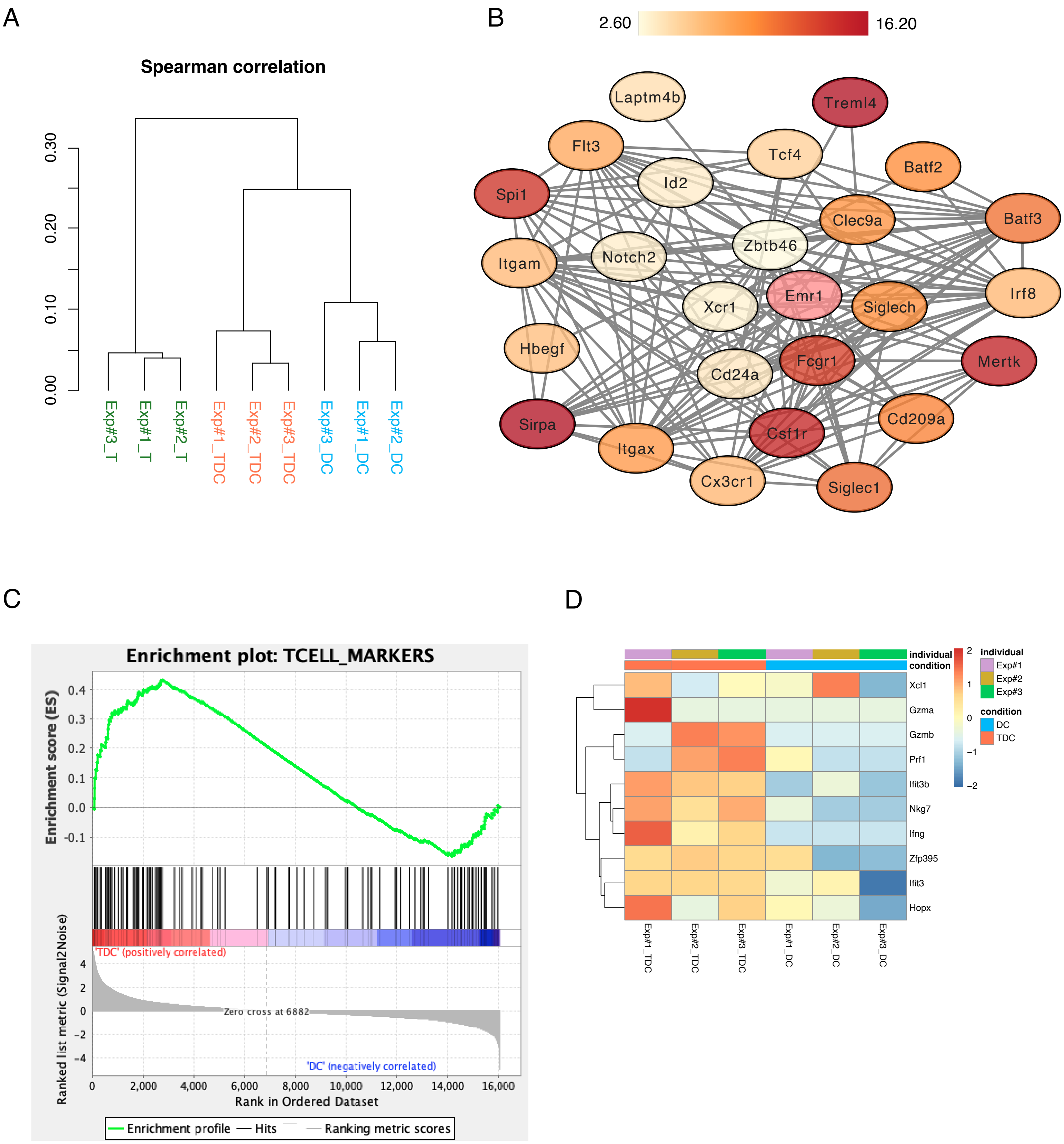

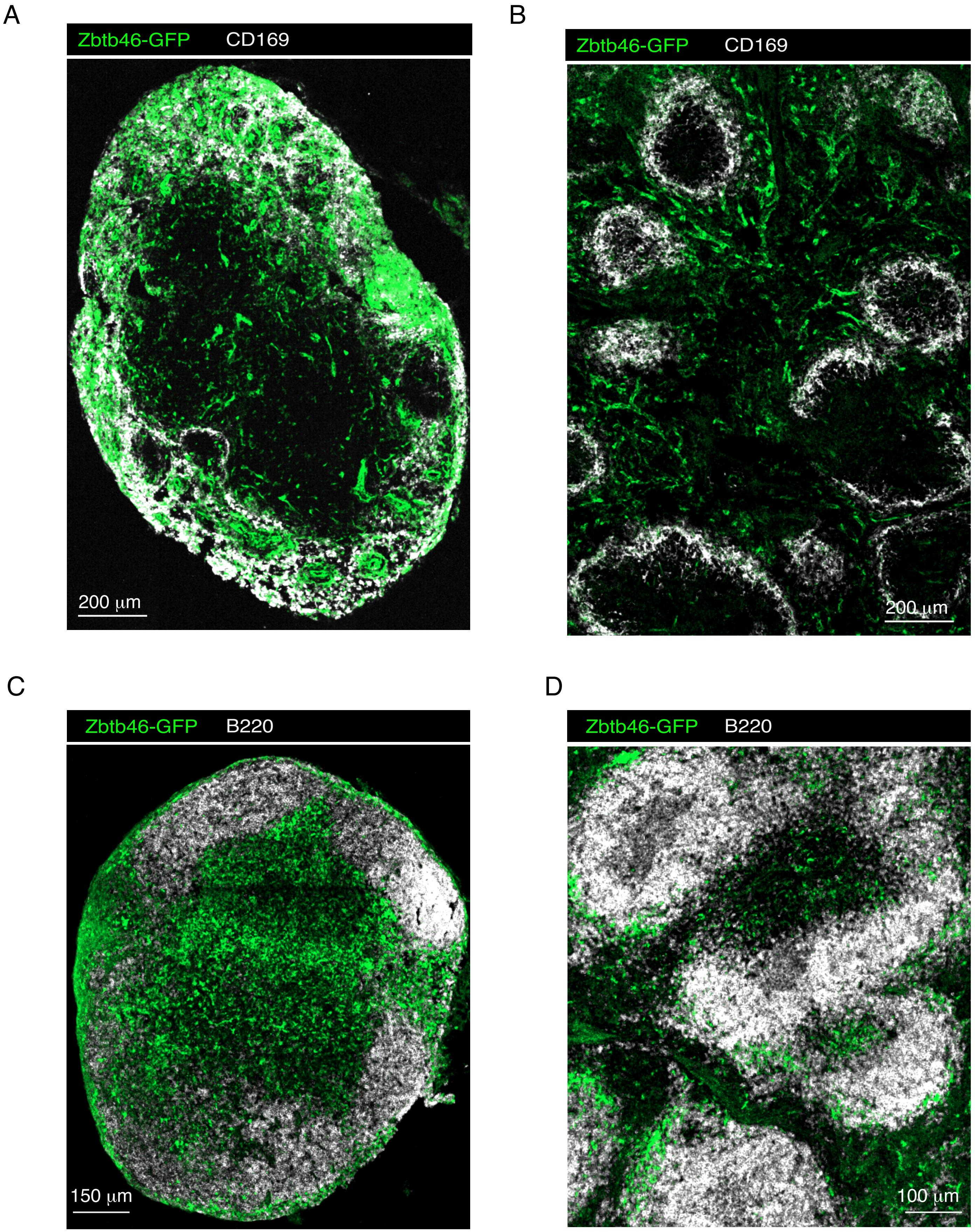

#### Figure S4

A

Popliteal LNs of naive mice

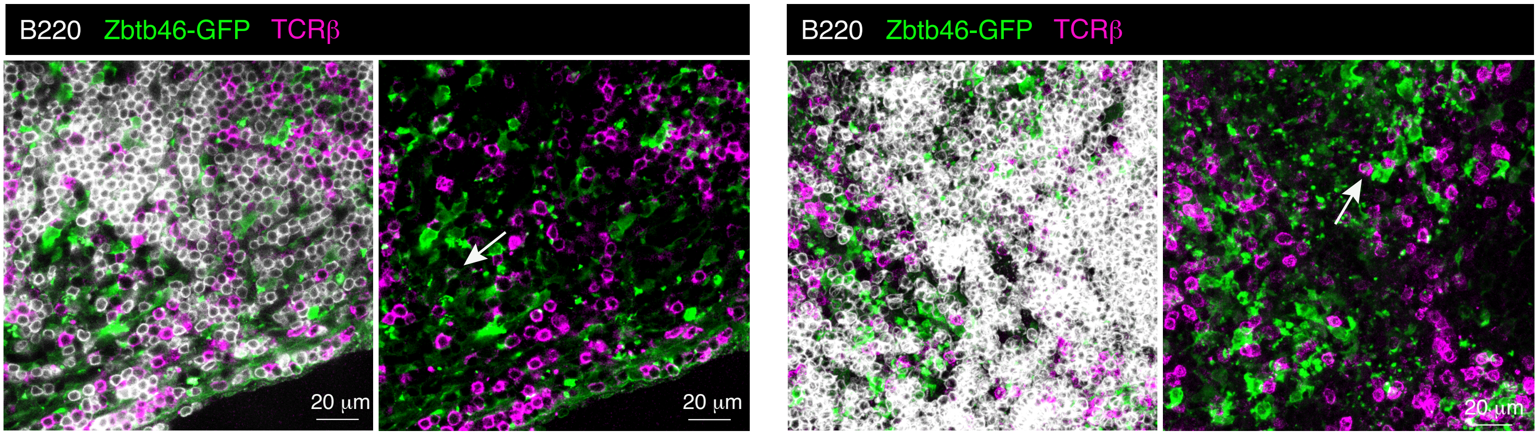


B

Spleens of naive mice

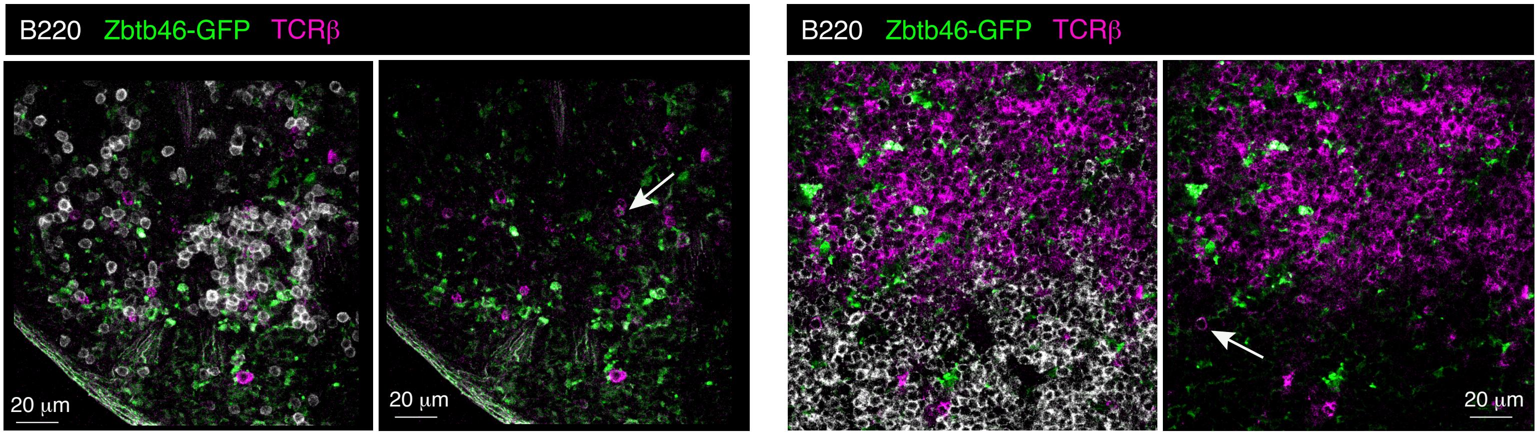

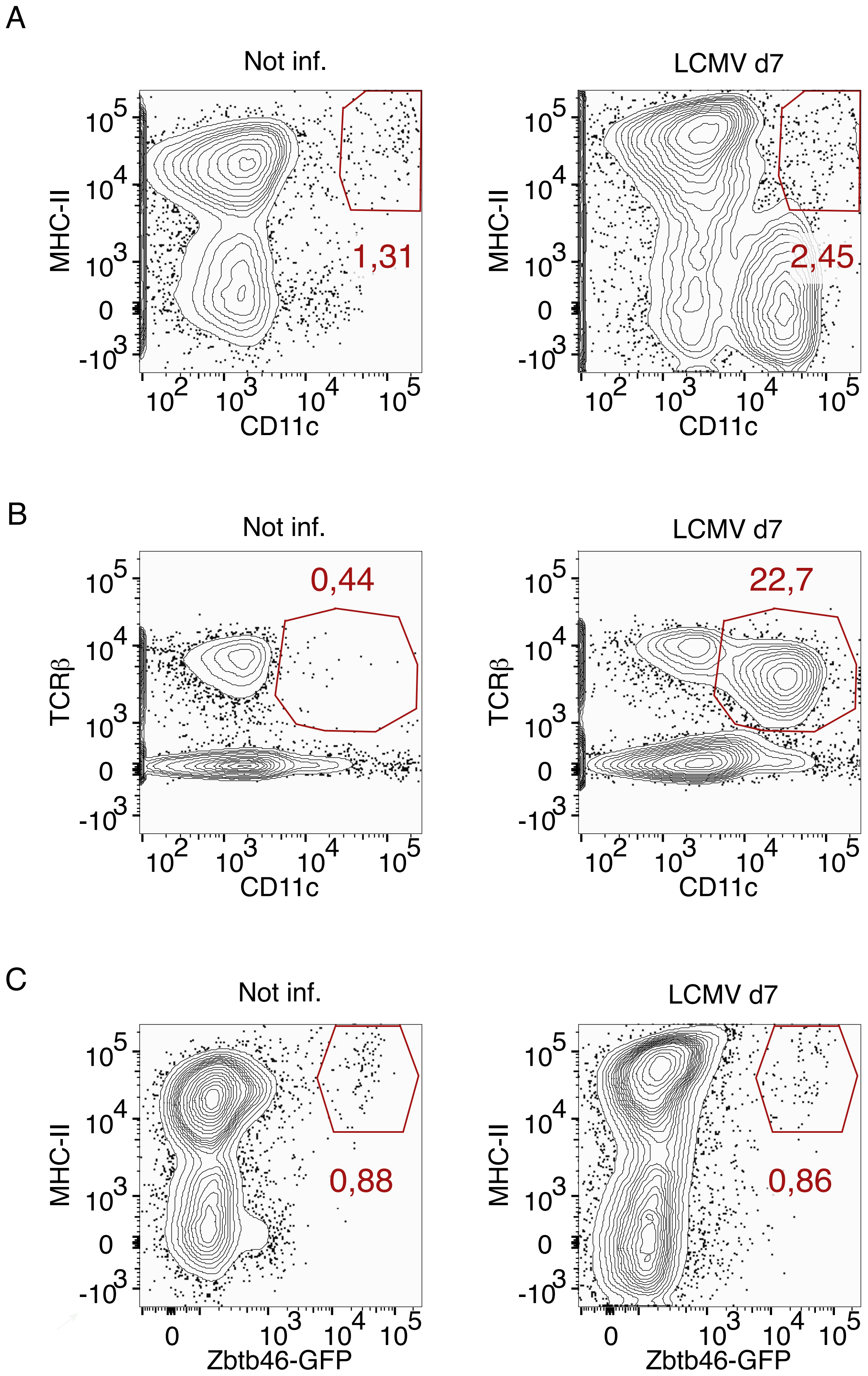

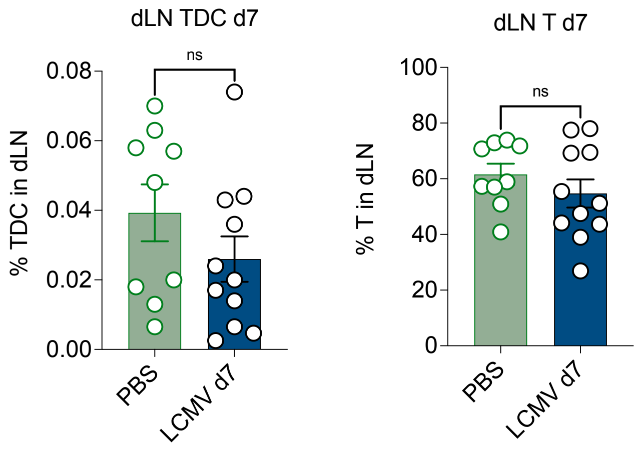

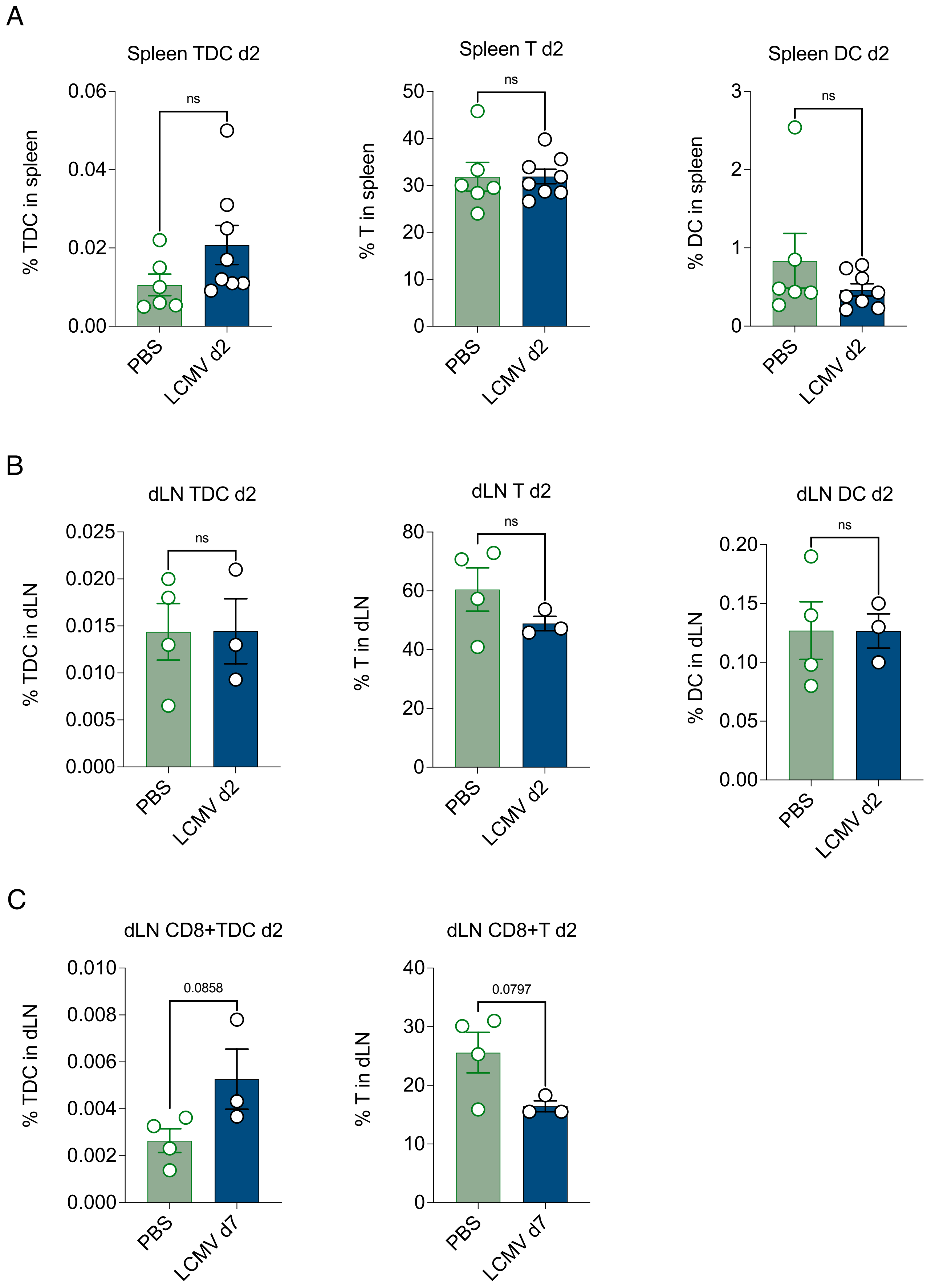

#### Figure S8

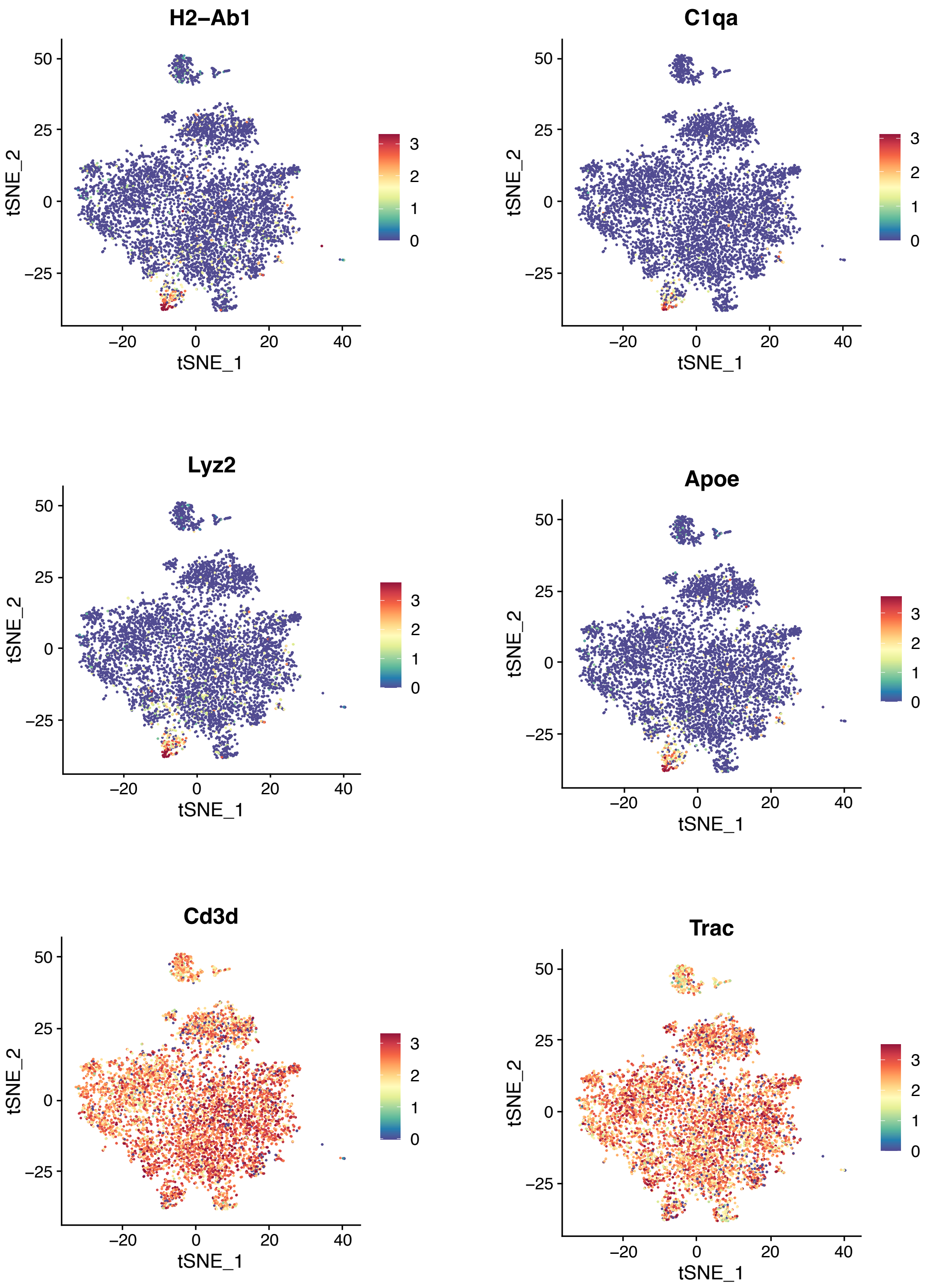

#### Figure S9

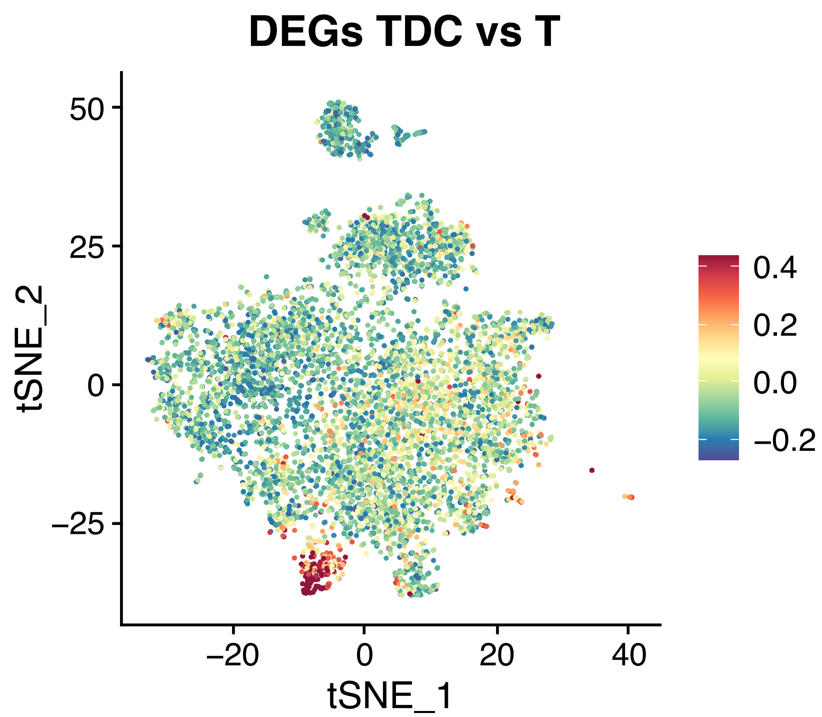
